# Smart squirrels use a mortise-tenon structure to fix nuts on understory twigs

**DOI:** 10.1101/2022.11.20.517261

**Authors:** Han Xu, Lian Xia, John R. Spence, Mingxian Lin, Chunyang Lu, Yanpeng Li, Jie Chen, Tushou Luo, Yide Li, Suqin Fang

## Abstract

Squirrels of temperate zones commonly store nuts or seeds under leaf litter, in hollow logs or even in holes in the ground; however, in humid rainforests, it is an evolutionary challenge for squirrels to hang elliptical or oblate nuts securely on trees to minimize germination or fungal infection. Here, we report a unique behaviour used by two species of small flying squirrels (*Hylopetes phayrei electilis* (G. M. Allen, 1925) and *Hylopetes alboniger* (Hodgson, 1870)) to cache nuts on Hainan Island. Squirrels intentionally carved grooves encircling ellipsoid nuts or distributed on the bottom of oblate nuts and used these grooves to fix nuts tightly between small twigs 0.1-0.6 cm in diameter and connected at angles of 25-40°. The resulting structures were similar to the mortise-tenon joint of ancient Chinese architecture. Cache sites were on small plants located 10-25 m away from the closest potentially nut-producing tree, a behaviour that likely deters discovery and consumption of the nuts by other animals. Their ability to shape individual nuts to store them more securely suggests it is a smart behavior, which is likely formed by the adaption to life in humid tropical rainforests to guarantee food supply and impacts the distribution of tree species.

## Introduction

Storing food is a common species-typical behaviour of squirrels and other rodents to prepare for seasons when nuts or seeds are in short supply (Andersson and Krebs, 1978; Steele et al., 2006). Nuts, in particular, are picked up from trees and then cached in various places. For example, many temperate-zone squirrels hoard nuts under leaf litter, or in holes in trees, logs or the ground (Cheng et al., 2005; Hadj-chikh et al., 1996). In subtropical zones, however, nuts may be stored by hanging them on tree branches, a behaviour thought to minimize germination or fungal infection in humid environments (Lichti et al., 2017; Xiao et al., 2013). These nuts are in clusters and are easily to be hanged on the branches directly, without needs to use tools to process nuts before being stored.

However, most nuts in tropical forests are single fruits with shapes that are mostly elliptical or oblate, something that makes them difficult to hang on trees. Solving this problem presents an evolutionary challenge to squirrels in such environments. The tool-use behaviors have been recorded for some animals in nature, such as chimpanzees, monkeys and corvids (Sanz et al., 2013). Squirrels also have smart decision-making behavior (Hadj-chikh et al., 1996; Hunt et al., 2021). Thus, we ask whether these nuts were processed by the squirrels’ tool using or some special behaviors to make them being securely fixed on the twigs.

In this paper, we show that a subspecies of the Indochinese Flying Squirrel, *Hylopetes phayrei electilis* (G. M. Allen,1925) and another species of the Flying Squirrel, *H. alboniger* (Hodgson, 1870) that co-occur in Hainan Island cache nuts by intentionally carving behavior before suspending them between the twigs of small plants. Both the two squirrel species are widespread in Southeast Asia (Duckworth et al., 2016; Tizard, 2016), but there is little published information about their habits and none from Hainan Province in China. This special behavior on nuts storage does not occur elsewhere in these two squirrels’ range and has not been reported that squirrels could carve the nuts intentionally to improve the success rate of storage. Thus, we set out to describe a unique behaviour through which they prepare and hang single ellipsoid or oblate nuts to solve the problem of providing safe storage in tropical rainforests. The sites where and the behavior how the nuts were stored and correlated squirrels’ activities were focused.

## Results and Discussion

A total of 151 cached nuts were found suspended on an overall total of more than 55 tree or shrub species distributed across 28 plant families in censuses of approximately 5.5 ha of forest **(Supplementary Figure 1, Supplementary Data file 1)**. Examples of nut locations and shaped nuts are shown in **Figures 1-2**. Most discovered nuts were fixed on a variety of small saplings and shrubs between plant twigs connected at angles of 25-40° **(Figure 3)**.

**Figure 1.**
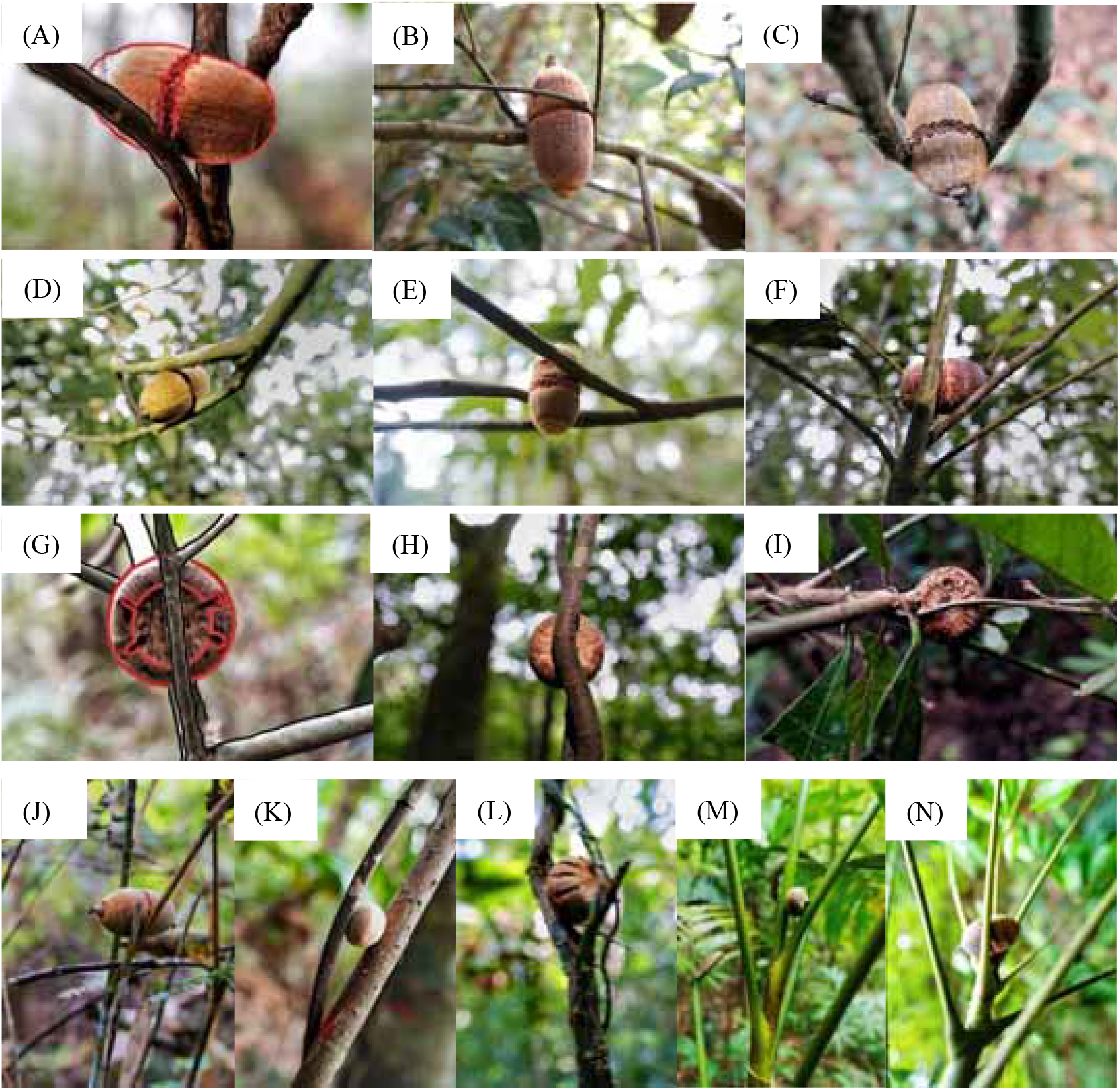
Nuts stored after surface preparation by flying squirrels. (**A**) Nut of *C. edithiae* (Skan) Schottky, with chewed grooves outlined in red. Nuts of *C. edithiae* fixed on trees, with (B-D) one, (E) two non-connected, or (F) spiral carved grooves encircling the nuts. (G) Nut of *C. patelliformis* (Chun) Y. C. Hsu et H. W. Jen, with chewed grooves outlined in red. (H-I) Nuts of *C. patelliformis* fixed on trees, with carved grooves on the bottom fixed on (J) bamboos and (K-L) lianas, between the big petioles of (M) trees and (N) palms.

**Figure 2.**
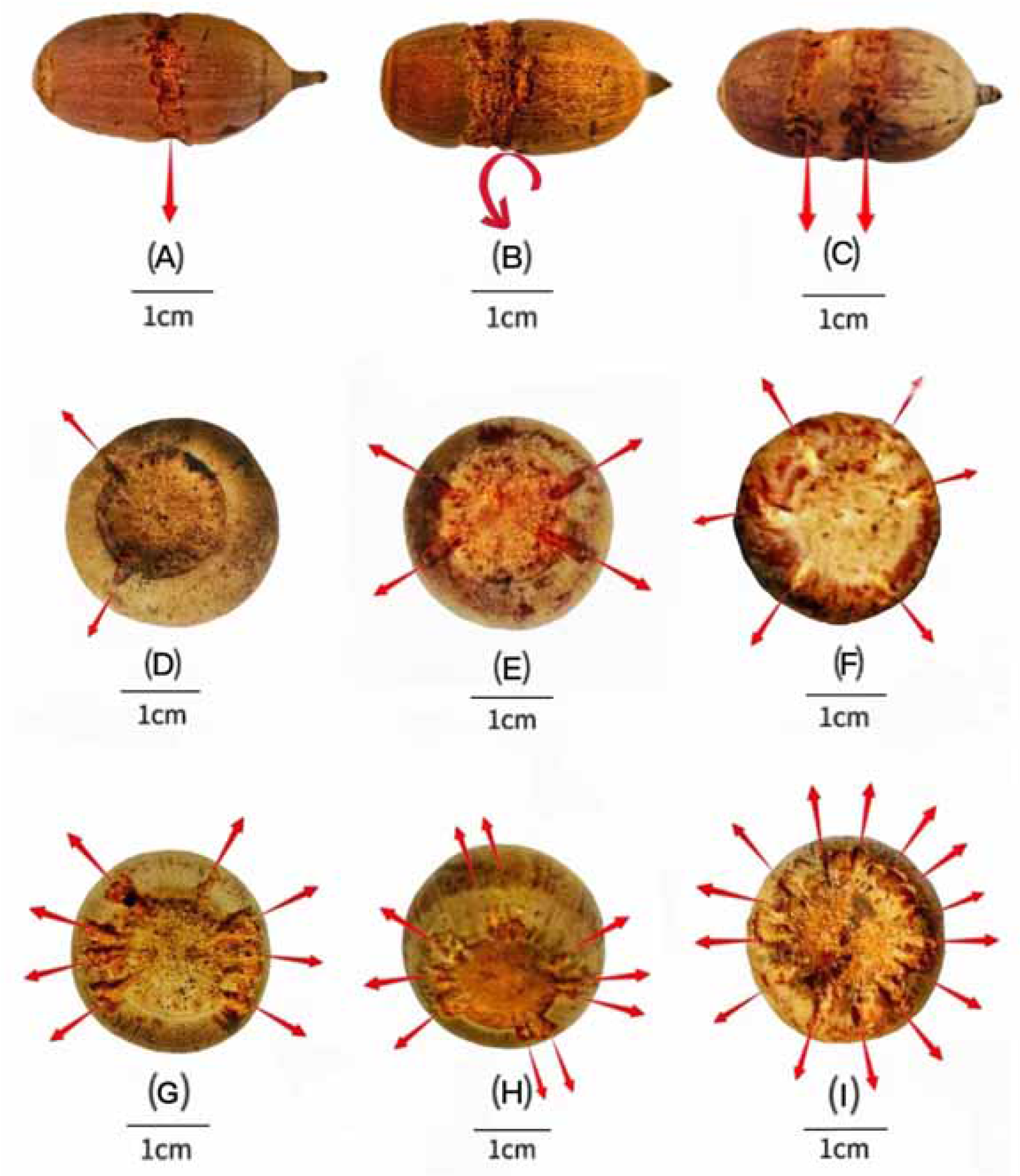
Variation in carved grooves depending on the storage situation. The carved surface grooves on nuts of *C. edithae* mostly encircle the middle of the nut, with (**A**) one, (B) one spiral, or (C) two separated grooves. The grooves on nuts of *C. patelliformis* are distributed on the bottom of the nuts, with (D) 2, (E) 4, (F) 6, (G) 8, (H) 10 symmetrically or (I) randomly distributed grooves.

**Figure 3.**
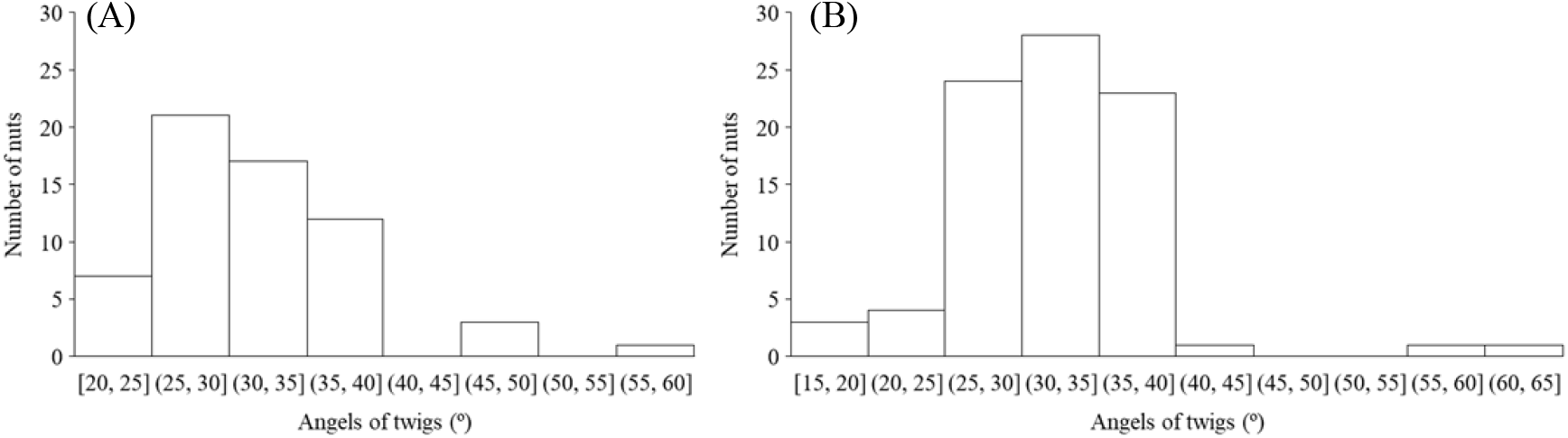
Nuts are fixed tightly between twigs generally meeting at angles of 25-40°. **(A)** *C. edithiae* nuts. (B) *C. patelliformis* nuts.

This range of angles accommodates the nut sizes of *Cyclobalanopsis edithiae* and *C. patelliformis* (2.4 cm (width) × 4.6 cm (length) and 2.4 cm (width) × 2.0 cm (height), respectively), which accounted for 96.7% of the nuts that we found cached (*C. edithiae* (40.4%), *C. patelliformis* (56.3%)). A few other nuts of *Lithocarpus fenzelianus* A. Camus and *C. fleuryi* (Hickel & A. Camus) Chun ex Q. F. Zheng were also found suspended on plants. Nuts of the two predominant species were disproportionately stored on small plants with diameters at breast height (DBH) of 0.4-1.6 cm **(Supplementary Figure 2)** and twig diameters of 0.10-0.60 cm **(Figures 4a, c)**. The diameters of the two twigs used for the storage structure were significantly correlated for *C. patelliformis* nuts but not for *C. edithiae* (*P*<0.001, **Figures 4b, d**). For nuts of *C. edithiae*, plant twig diameter was significantly correlated with the nut groove width, and generally varied from 0.20 cm to 0.60 cm (*P*<0.001, **Figure 5**). The first to third branches at a plant height of 1.50-2.50 m comprised 45.9% of *C. edithiae* and 43.5% of *C. patelliformis* storage sites we discovered **(Supplementary Figure 3)**.

**Figure 4.**
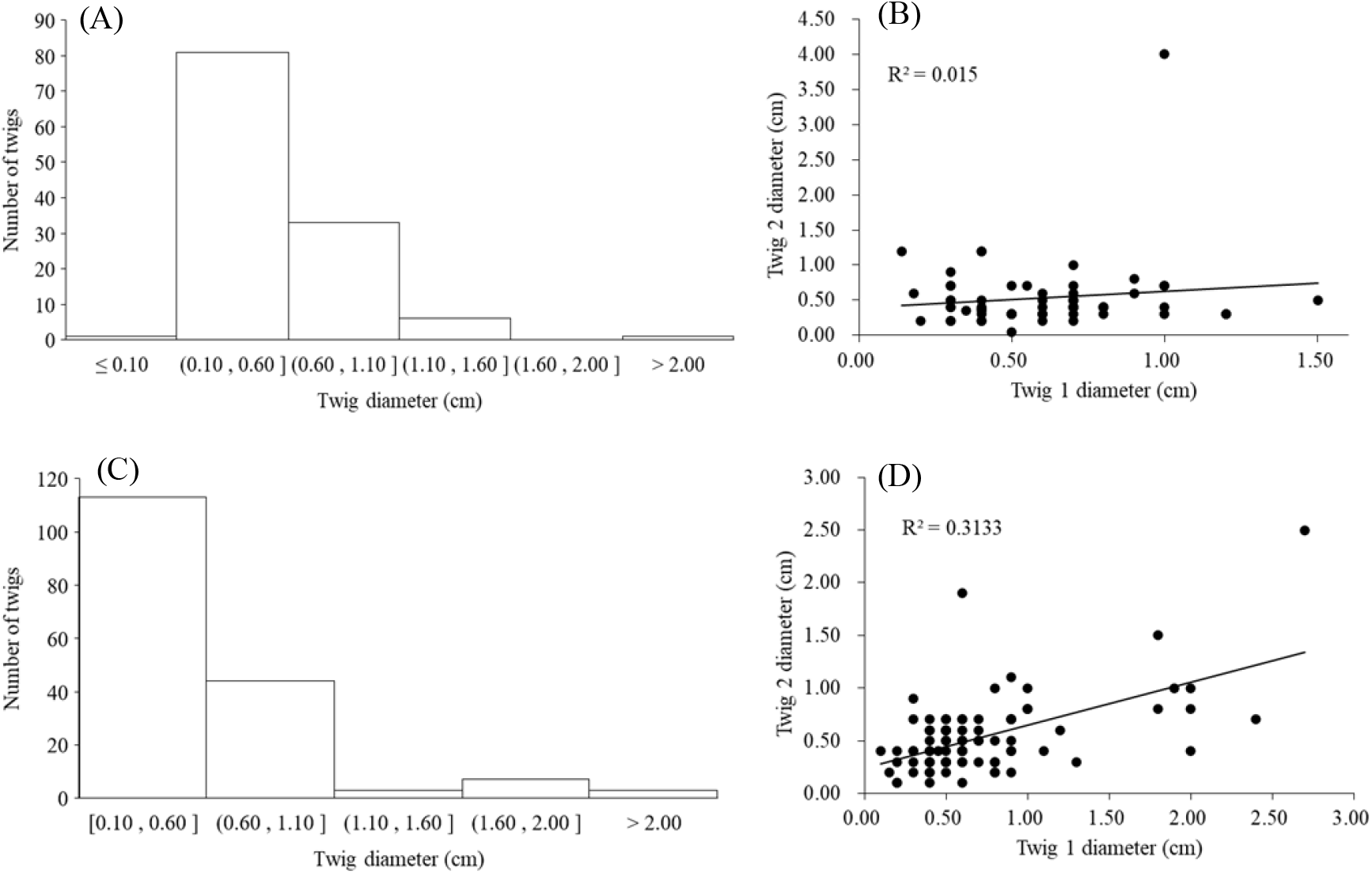
Nuts were stored mainly on small plants between twigs with diameters of 0.10 - 0.60 cm. (**A**) Histogram of diameters of twigs used to store nuts of *C. edithiae*. (B) Diameters of twigs used to store *C. edithiae* nuts were not significantly correlated. (C) Histogram of diameters of twigs used to store nuts of *C. patelliformis*. (D) Diameters of twigs used to store *C. patelliformis* were significantly correlated. Twig 1 and twig 2 are two twigs fixed the nuts.

**Figure 5.**
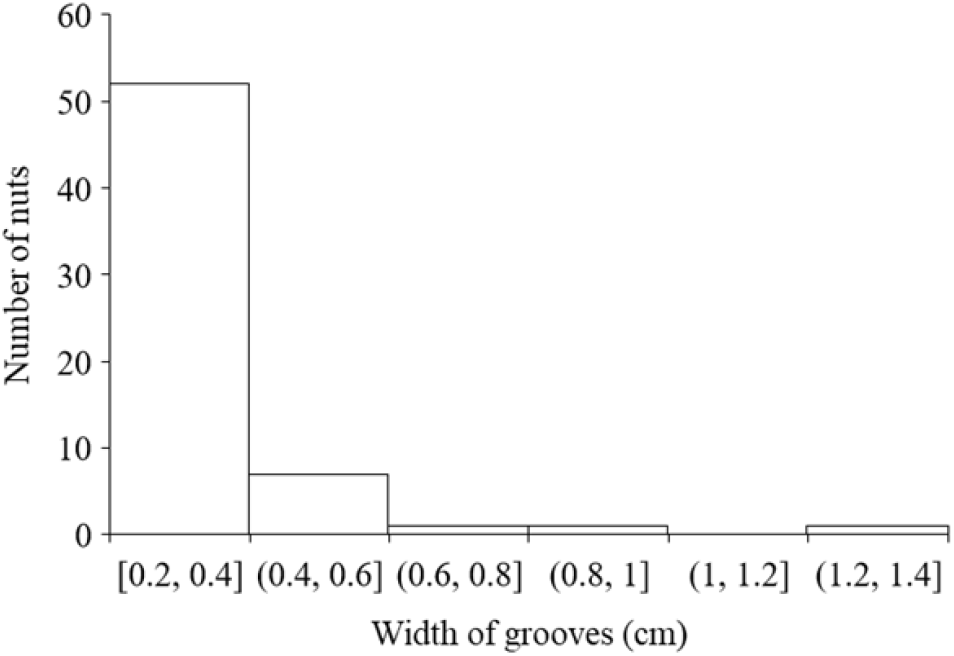
Grooves carved by squirrels on most *C. edithiae nuts* were 0.2 - 0.6 cm in width.

We used the footage from our infrared cameras to identify the nocturnal flying squirrels, *Hylopetes phayrei electilis* and *H. alboniger*, as being actively associated with these nuts **(Supplementary Media files 1-5)**. Both *H. phayrei electilis* and *H. alboniger* are small-bodied flying squirrels found in tropical forests from Myanmar, south to northwestern Vietnam and east into the Guizhou, Guangxi and Fujian Provinces of southern China. These two species co-occur in the tropical mountain rainforests in the Hainan, Province of China (Li et al., 2012).

Squirrels of both species apparently carve spiral zigzag surface grooves that encircle the midsection of the ellipsoid nuts of *C. edithiae* **(Figures 1a-f)** with one or occasionally two grooves **(Figures 2a-c)**. The two non-connected or spiral grooves appear to be useful for adjusting the storage of nuts to the specific orientation of the twigs. In contrast, up to 20 grooves are carved on the bottom of the oblate nuts of *C. patelliformis* **(Figures 1g-i)**, commonly with 0-8 shallow scattered grooves **(Supplementary Figure 4a)**, some even have 2, 4, 6, 8, 10 symmetric grooves **(Figures 2d-i)**. These oblate nuts stored on living trees and shrubs have significantly more carved shallow scattered grooves than those stored on dead trees and lianas (5.1±5.0 *vs*. 2.8±4.0, t=2.1591, df=46.402, *P*=0.036). The bark of dead trees and lianas is coarser than that of living trees, and thus dead stems may require fewer grooves to hold the nuts securely. We also note that that the grooves on the ellipsoid nuts of *C. edithiae* are deeper (more than 0.5 mm) than on the oblate nuts of *C. patelliformis* (less than 0.45 mm, **Supplementary Figure 4b**), but grooves of either depth do not damage the endosperm of nuts, thus likely reducing potential impacts of fungi during the storage period.

Surface preparations by the squirrels allow the nuts to be “pressure fitted” between the two plant twigs in a way similar to the mortise-tenon structure in ancient Chinese architecture (Qiao et al., 2021) **(Figure 2)**. The carved nuts are inlayed between plant twigs (0.10-0.60 cm in diameter) intersecting at specific angles (25-40°) on various understory plants **(Figure 3)**, including small trees and shrubs, sometimes lianas, bamboos or dead trees, and even occasionally big petioles of palms or trees **(Figures 1j-n, Supplementary Table 2)**. Once fixed in this manner nuts are resistant to being blown off by strong wind or shaking that we administered **(Supplementary Media files 6-11)**.

We found that nuts produced by large trees of the two *Cyclobalanopsis* species are stored on much smaller understory plants by these flying squirrels. The distance between nut producing trees and storage sites varied from 10 to 25 m **(Supplementary Figure 5)**, distances larger than the average canopy width of large trees that we have observed in the Jianfengling forest. This likely reduces discovery by other animals searching for aboveground nuts below the parent trees (Cao et al., 2011), although it seems that some nuts may be still eaten by mice. Based on our assessments, only 63.6% nuts stored on understory plants were fresh at the time of survey, meaning that they can persist in these storage sites for lengthy periods. Over the 44 days between the first and second surveys, 19.7% of the stored nuts had disappeared, and 15.0% of the nuts discovered during the second survey were new. Over the 61 days between the second and third surveys, 43.7% of the stored nuts had disappeared, and 20.6% of the nuts discovered were new. Thus, in a general sense, the number of nuts being stored seems to decrease gradually from January to May after the fruiting season.

Seedlings that result from any dropped, fallen or forgotten nuts **(Supplementary Figure 6)** may germinate at some distance from their parents. In this way, seed dispersal by these squirrels may decrease competition between seedlings and parent trees and thus increase seedling survival rates that could, in turn decrease the negative density dependence of conspecific trees as consistent with recently published arguments (Detto et al., 2019). Unfortunately, we are presently unable to estimate what proportion of the disappearance results from use by squirrels, although our video footage establishes that some nuts are indeed removed **(Supplementary Media files 1-5)**. Nonetheless, some proportion of nuts likely falls and may germinate near the storage site, as is common for seeds and nuts cached by squirrels, especially in hardwood forests (Steele and Yi, 2020). Furthermore, although the importance of large DBH trees has been emphasized for the maintenance of forest ecosystem productivity (Lutz et al., 2018), from a broader perspective, small understory plants may help sustain the diversity and complexity of forest structure in the long run through plant-animal interaction processes like those described here.

The precipitation and the humidity of the environments may be the motivation to drive squirrels to fix the nuts above the ground. In the temperate-zone, the annual precipitation is usually less than 1000 mm and the coniferous trees are dominant, which produce relatively dry fallen needles. The nuts could be safely stored under leaf litter or on the ground without special processing (Hadj-chikh et al., 1996). In the studied sites in the tropical forests in Jianfengling, Hainan Island, the annual precipitation varies from 1300-3700 mm (Xu et al., 2015). The nuts may germinate or been infected by fungi in the humid environments, especially in the rainy season. Hence, it is much better for the squirrels to store nuts above the ground, such as on the understory twigs.

For the special nuts carving behavior, we guess it may be originated from an inadvertent carving attempt long long ago and squirrels learned that the carved grooves help improve the nut fixation between the twigs. This learning ability is widely reported in animals, e.g., wild parrots have strong abilities to use and manufacture tools, and imitate (Shaw, 2021). Gradually, the special nuts carving behaviour became a common practice for these two squirrels, although this speculation needs more evidences like animal cranial zootomy or behaviour training experiments in the future.

In summary, individuals of *H. phayrei electilis* and *H. alboniger*, among the nine species of squirrels known from tropical forests of Hainan **(Supplementary Table 3)**, store nuts of different shapes and sizes securely on a variety of plant twigs at some distance from the plants that produced them. Clearly, squirrels have evolved the ability to prepare the nuts for such storage by carving grooves in their surfaces to create a “mortise-tenon” connection between nuts and plant twigs selected for particular characteristics. This leads to the inference that squirrel can reason about how to effectively reshape the nuts for storage. The symmetric groove structure on oblate nuts carved by individuals of *H. phayrei electilis* and *H. alboniger* support a complex and effective strategy for fixing nuts.

Taken together, these observations indicate that these two flying squirrels have adapted to the particular humid tropical rainforest circumstances and developed smart behavior flexibly to solve nut-storage problems. We suggest that this is a smart behavior that has evolved through plant-animal-environment interactions that provide long-term adaption to secure food, given the high precipitation environment and the seasonal production of nuts before the coldest month in these rainforests. The caching behavior may further affect dispersal and the functional traits of nuts (Chang and Zhang, 2014; Xiao et al., 2004) in a way that alters the spatial and temporal distribution pattern of the local plant community in the long-run. These possibilities deserve more attention in the future (Goheen and Swihart, 2003; Rong et al., 2013).

## Materials and Methods

### Study site

This study was conducted in the Jianfengling region of Hainan Tropical Rainforest National Park in Hainan Province, China (108°46’-109°45’E). The area has a seasonal tropical monsoon climate with a rainy season from June to October and a dry season from November through May of the next year. The mean annual temperature is 19.7°C and the annual average precipitation is 2684 mm. This is the second rainiest area on Hainan Island, with an average annual relative humidity over > 88%. The forests include 992 free-standing tree and shrub species, and are dominated by trees of Fagaceae, Lauraceae and Moraceae (Xu et al., 2012). *Castanopsis, Lithocarpus* and *Cyclobalanopsis* are three main genera of Fagaceae, which reproduce nuts which are used as food by mammals. Cupules of *Castanopsis* are solitary on rachis, completely or partially enclosing the nut, cupules of *Lithocarpus* are grouped together in cymes on rachis, completely or partly enclosing the nut, while cupules of *Cyclobalanopsis* are solitary, and do not enclose the nuts. The enclosed nuts are not easy for squirrels to deal with. Thus, *Cyclobalanopsis* nuts are highly preferred as food by squirrels or other animals. *Cyclobalanopsis edithiae (Skan) Schottky* and *Cyclobalanopsis patelliformis* (Chun) Y. C. Hsu et H. W. Jen are the two most abundant species that have naked nuts in the mountain forests of Jianfengling (Xu et al., 2015) **(Supplementary Table 1)**. Both are in fruit from October to December, just before the coldest month (January) in Hainan.

### Field investigation

This work was prompted by inadvertent discovery of *Cyclobalanopsis* nuts with strange surface grooves tucked into the Y-shaped crotches of twigs on understory plants **(Figure 1)**. Most of the regular grooves showed signs of having been chewed by some animal(s) **(Figure 2)**, perhaps in order to increase the friction between nuts and plant twigs so as to fix the nuts securely in place. Thus, we conducted a systematic field investigation from January to May 2022 to discover the animal(s) involved and to study the nut caching behavior in more detail.

We first made a systematic search of a ca. 5.5 ha forest area for grooved nuts suspended in vegetation to establish the relationship of this phenomenon to various plant types. The species identity, diameter at breast height (DBH), geographical position, and elevation of plants found with chewed nuts were recorded. We also measured the diameter and angles of the two twigs where the nuts were fixed. Local stand canopy closure was estimated visually and the dominant trees within a ca. 20 m radius of the stored nuts were recorded. Finally, we measured the distance between the storage site and the nearest tree where the nuts could have been produced. For the suspended nuts themselves, we determined the species, weight, diameter and height on each plant. The number, depth and position of the surface grooves were also noted. Because precise measurement was impossible, depth of the carved grooves was measured as a categorical variable classified in the following three groups: shallow (ca. 0-0.15 mm), medium (ca. 0.15-0.30 mm) and deep (ca. 0.30-0.45 mm). We also recorded whether each nut was fresh, eaten by insects or infected by fungi; fresh nuts are green but become wrinkled and black with age. All discovered nuts were photographed.

The first search for cached nuts was carried out on January 15, 2022. We re-surveyed the site 44 days later on February 28 to learn whether the nuts recorded previously were still where we initially found them, and to find any new nuts that had been stored in the area. A third survey was made 61 days later on April 30.

Because we initially knew neither the identity of animals that stored the nuts nor how nuts were stored and retrieved, we set up 32 motion-activated infrared cameras (22 WildINSights 20MP 1080P HD Trail Cameras and 10 WildINSights 5MP 960P HD Trail Color Cameras) to monitor animal activities related to storage or consumption of the nuts. These cameras were positioned to view typical nuts that we found and their surroundings. Animals filmed as being associated with the nuts were subsequently identified to species by experts using the resulting pictures and videos.

### Statistical analyses

Histograms were drawn to describe variation in the sizes and location of storage plants and nuts. Standard *t*-tests were used to assess the significance of differences between paired sets of variables and linear regression was used to assess relationships between sizes of the two twigs that constituted the nut storage site.

## Acknowledgments

We thank Wenhao Qin, Chuanwen Yu, Fenglin Huang and Tao Zhang for helping to collect the field data. We also appreciate suggestions from Yi Ding from Chinese Academy of Forestry that improve the manuscript, and assistance from Fang Liu from Chinese Academy of Forestry and Qiang Zhang from Institute of Zoology, Guangdong Academy of Sciences in identifying the animals.

## Data availability

All data are available in the main text or the supplementary files.

## Supplementary Files

**Supplementary Figures 1 to 6**

**Supplementary Tables 1 to 3**

**Supplementary Media files 1 to 5.**

**Footage from infrared cameras of squirrels checking and removing nuts from the storage sites**.

There are two folders, each with footage about one of the two squirrel species. In the “By *Hylopetes alboniger*” folder, there are three media files about this species, including

Media 1, shows squirrel 1 checking nuts at the storage sites.

Media 2, shows squirrel 2 removing nuts from storage sites.

Media 3, shows squirrel 3 removing nuts from storage sites.

In the “By *Hylopetes phayrei electilis*” folder, there are two media files about this species, including

Media 4, shows squirrel 4 checking nuts at storage sites.

Media 5, shows squirrel 5 removing nuts from storage sites.

**Supplementary Media files 6 to 11**.

**Footage showing that shaking the plants does not dislodge nuts stored by squirrels**.

There are two folders, one for each of the two principal Fagaceae species.

In the “*Cyclobalanopsis edithiae*” folder, there are three media files, including

Media 6, shows footage of shaking a liana with a stored nut.

Media 7, shows footage of shaking a sapling with a stored nut.

Media 8, shows footage of shaking a sapling with a stored nut.

In the “*Cyclobalanopsis patelliformis*” folder, there are three media files, including

Media 9, footage of shaking a liana with a stored nut.

Media 10, footage of shaking a sapling with a stored nut.

Media 11, footage of shaking a sapling with a stored nut.

## Supplementary Data file 1

**The plants used to store nuts and their growth form**.

**Supplementary Figure 1.**
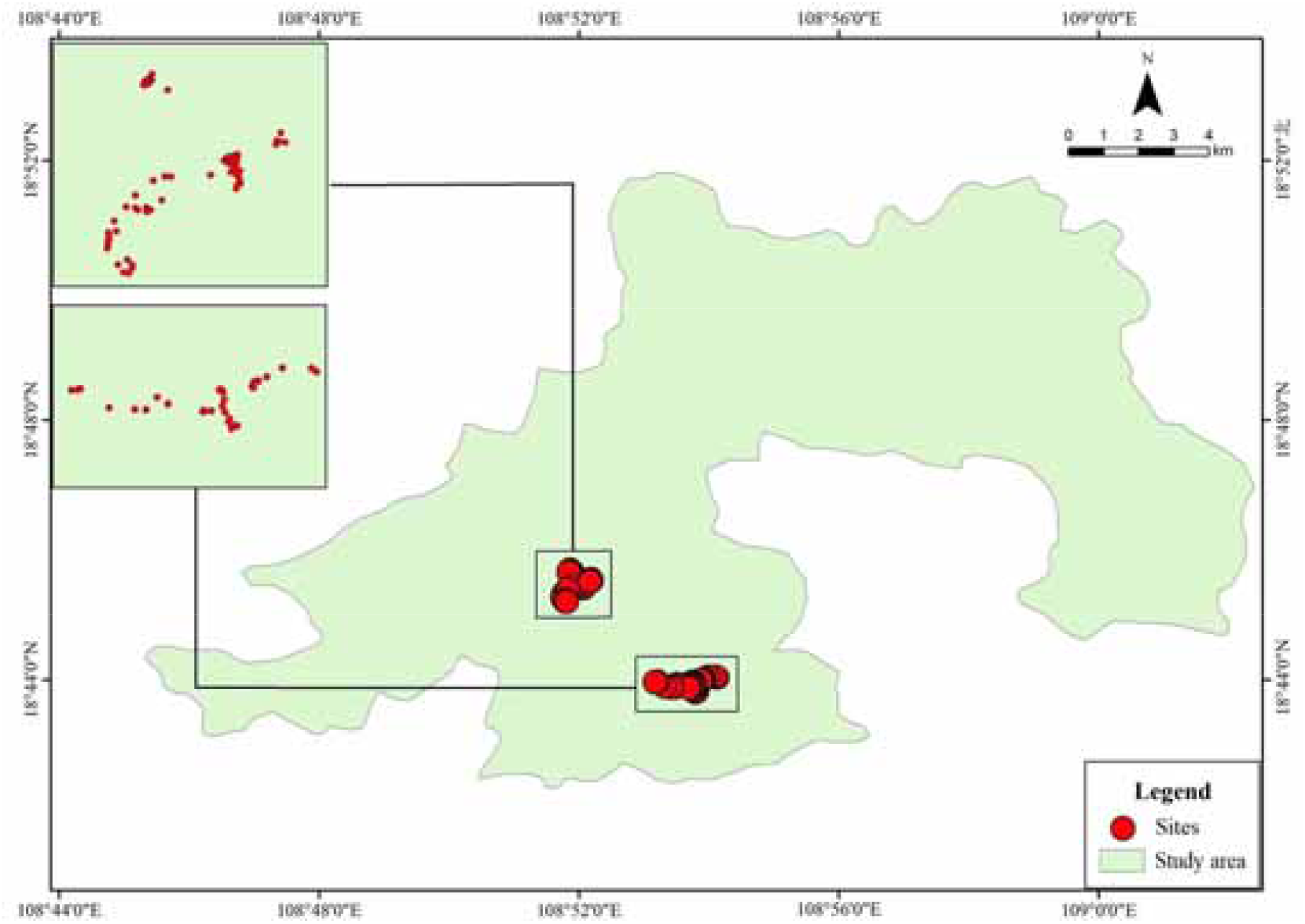
Spatial distribution of the 151 suspended nuts observed in Jianfengling Nature Reserve, Hainan, China.

**Supplementary Figure 2.**
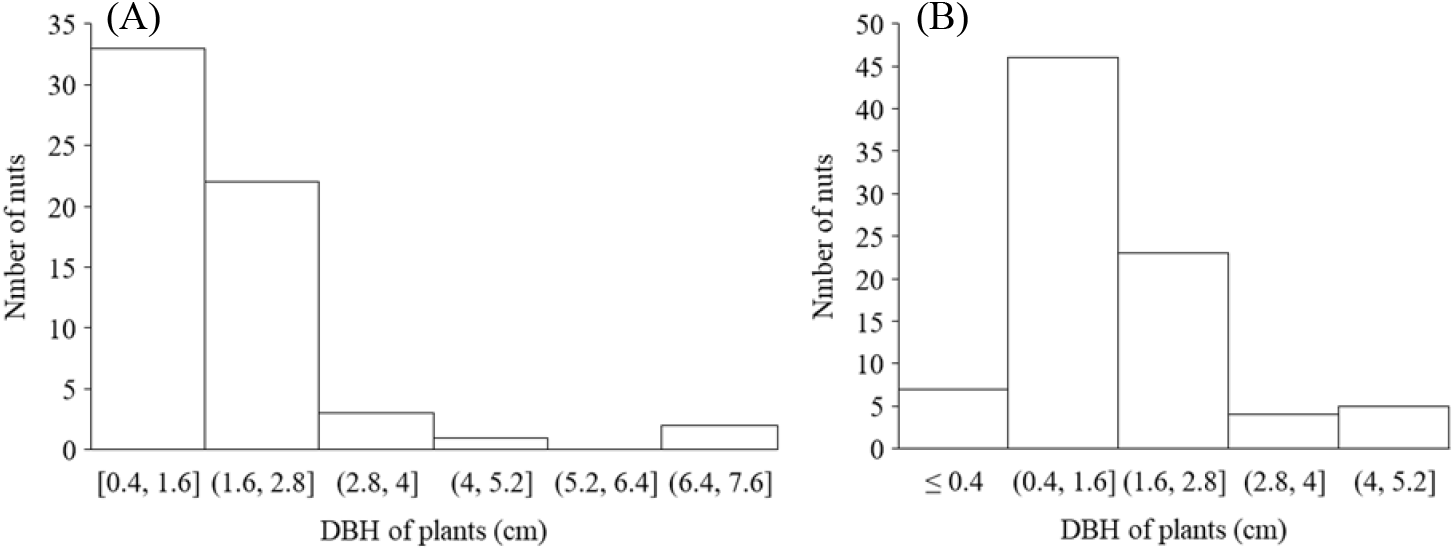
Most nuts were stored on small plants with diameter at breast height (DBH) ranging from 0.4 - 1.6 cm. (A) *C. edithiae* nuts. (B) *C. patelliformis* nuts.

**Supplementary Figure 3.**
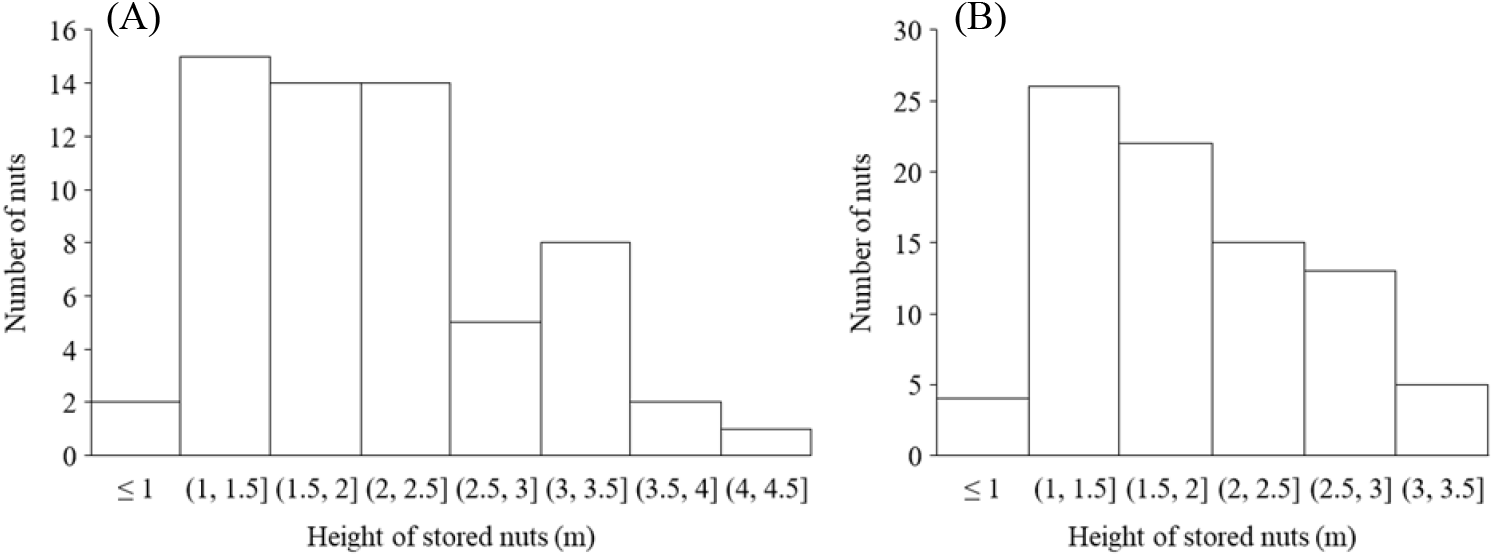
Nuts were generally stored on the first to third branches at 1.5-2.5 m aboveground. (A) *C. edithiae* nuts. (B) *C. patelliformis* nuts.

**Supplementary Figure 4.**
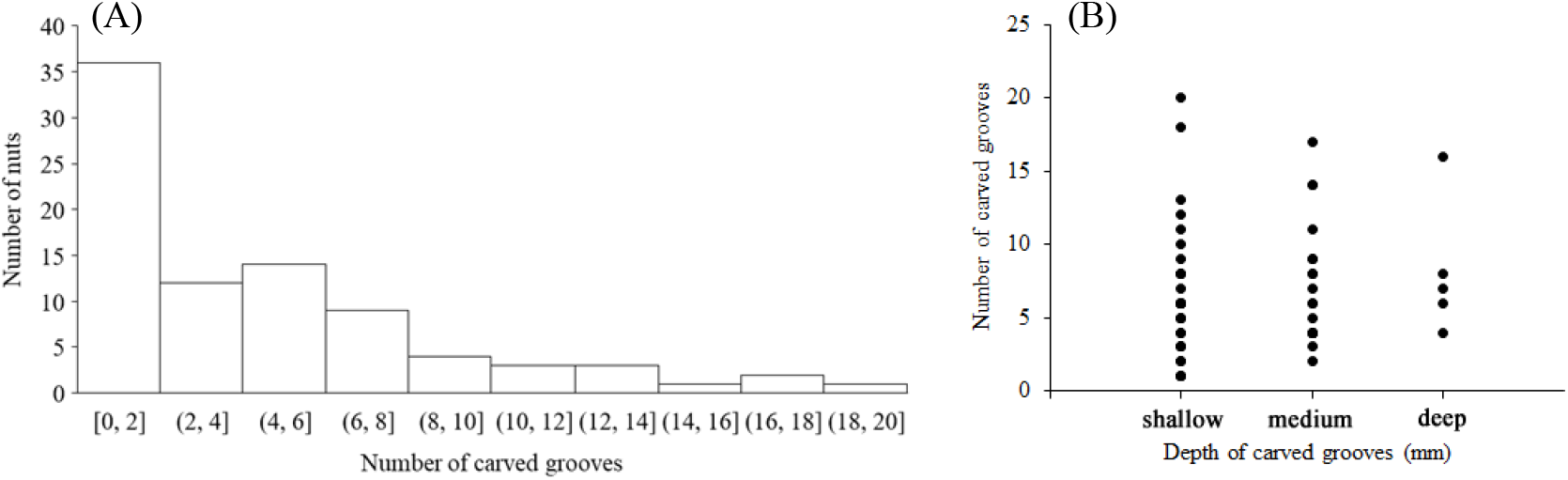
Number of grooves carved on the oblate nuts of *C. patelliformis*. (A) Most nuts had fewer than 8 grooves. (B) The depth of most grooves was shallow to medium.

**Supplementary Figure 5.**
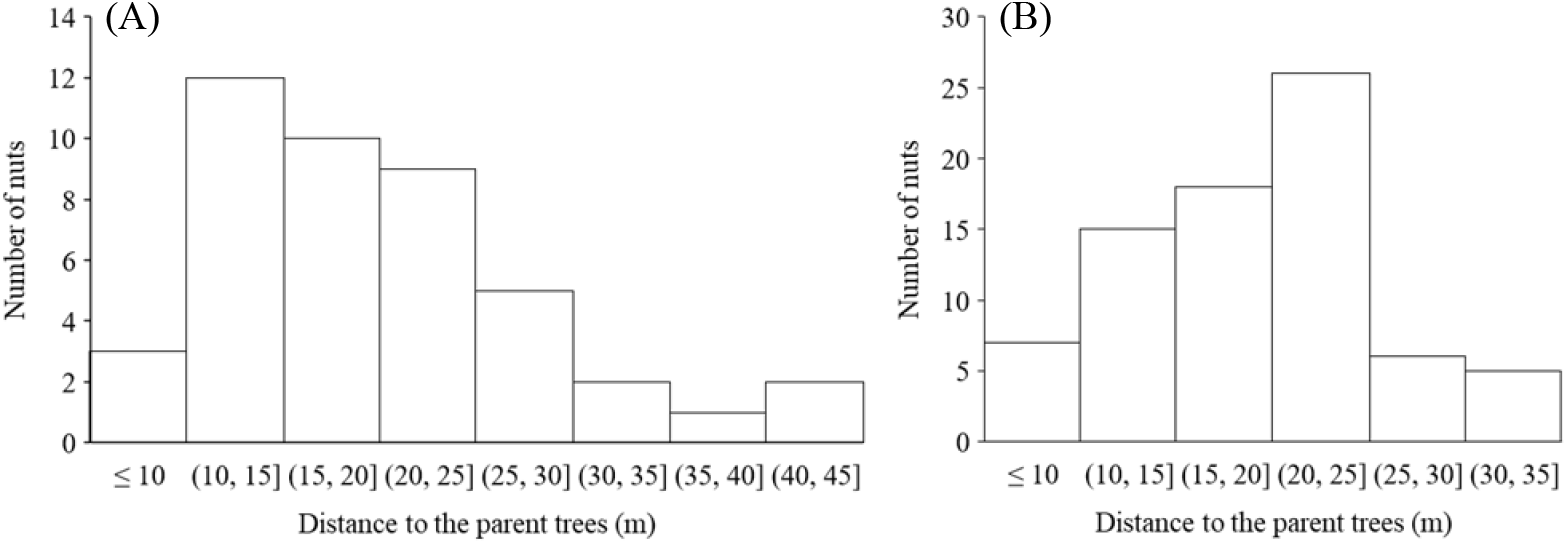
Distance from storage sites to potential parent trees for the nuts varied from 10 m to 25 m. (A) *C. edithiae* nuts. (B) *C. patelliformis* nuts.

**Supplementary Figure 6.**
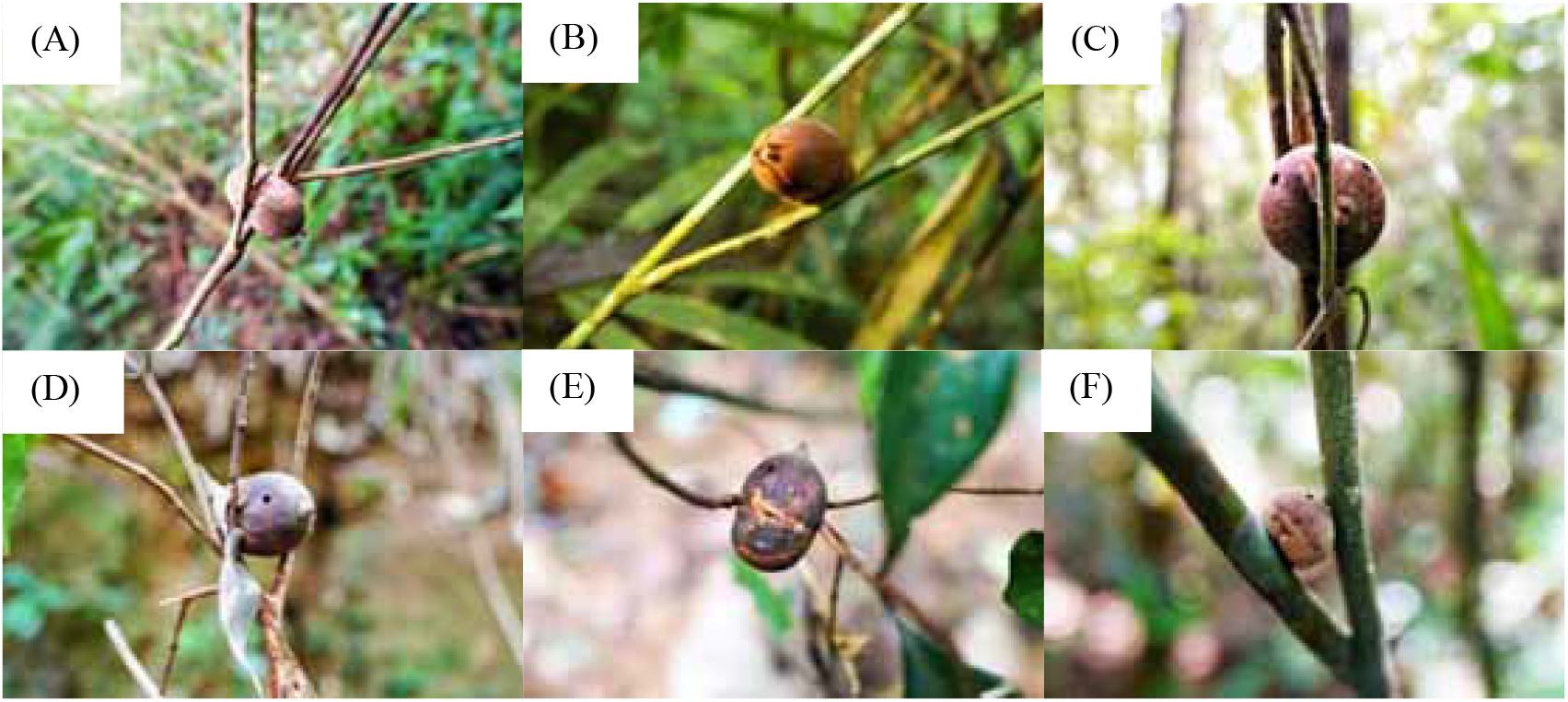
After long (e.g., >ca. 365 days) storage, nuts become not fresh. (A) lost germination ability, (B) germinated or (C-E) were destroyed by insects.

**Supplementary Table 1.**
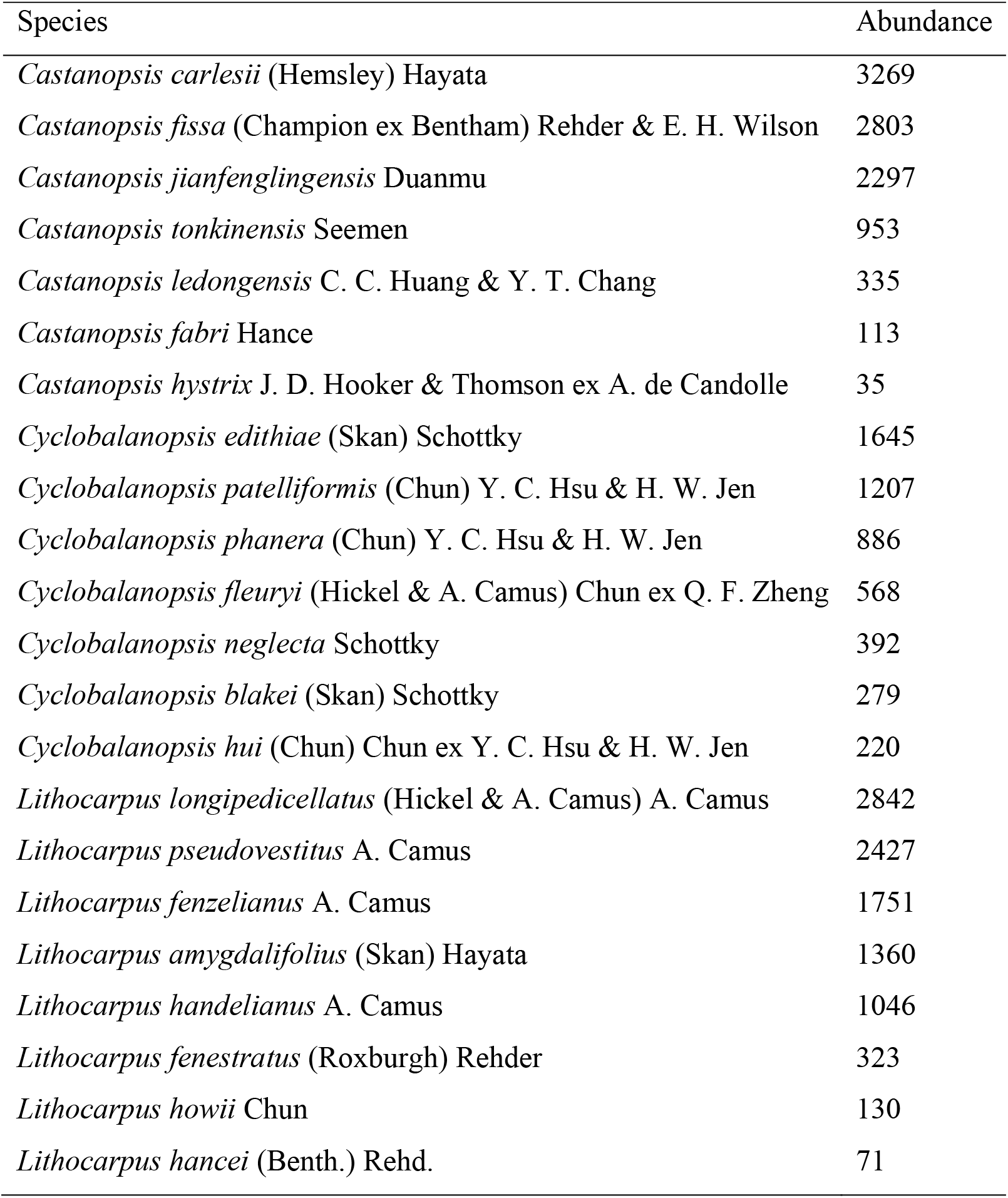
Main Fagaceae species found in a 60 ha plot in the Jianfengling forests.

| Species | Abundance |
| --- | --- |
| *Castanopsis carlesii* (Hemsley) Hayata | 3269 |
| *Castanopsis fissa* (Champion ex Benth) Rehder & E. H. Wilson | 2803 |
| *Castanopsis jianfenglingensis* Duanmu | 2297 |
| *Castanopsis tonkinensis* Seemen | 953 |
| *Castanopsis ledongensis* C. C. Huang & Y. T. Chang | 335 |
| *Castanopsis fabri* Hance | 113 |
| *Castanopsis hystrix* J. D. Hooker & Thomson ex A. de Candolle | 35 |
| *Cyclobalanopsis edithiae* (Skan) Schottky | 1645 |
| *Cyclobalanopsis patelliformis* (Chun) Y. C. Hsu & H. W. Jen | 1207 |
| *Cyclobalanopsis phanera* (Chun) Y. C. Hsu & H. W. Jen | 886 |
| *Cyclobalanopsis fleuryi* (Hickel & A. Camus) Chun ex Q. F. Zheng | 568 |
| *Cyclobalanopsis neglecta* Schottky | 392 |
| *Cyclobalanopsis blakei* (Skan) Schottky | 279 |
| *Cyclobalanopsis hui* (Chun) Chun ex Y. C. Hsu & H. W. Jen | 220 |
| *Lithocarpus longipedicellatus* (Hickel & A. Camus) A. Camus | 2842 |
| *Lithocarpus pseudovestitus* A. Camus | 2427 |
| *Lithocarpus fenzelianus* A. Camus | 1751 |
| *Lithocarpus amygdalifolius* (Skan) Hayata | 1360 |
| *Lithocarpus handelianus* A. Camus | 1046 |
| *Lithocarpus fenestratus* (Roxburgh) Rehder | 323 |
| *Lithocarpus howii* Chun | 130 |
| *Lithocarpus hancei* (Benth.) Rehd. | 71 |

**Supplementary Table 2.** The types of plants used for nut storage.

| Plant type | Number of individuals | Percentage of all individuals (%) |
| --- | --- | --- |
| Alive tree | 108 | 71.5 |
| Dead tree | 17 | 11.3 |
| Alive liana | 19 | 12.6 |
| Dead liana | 2 | 1.3 |
| Bamboo | 5 | 3.3 |
| Total | 151 | 100 |

**Supplementary Table 3.**
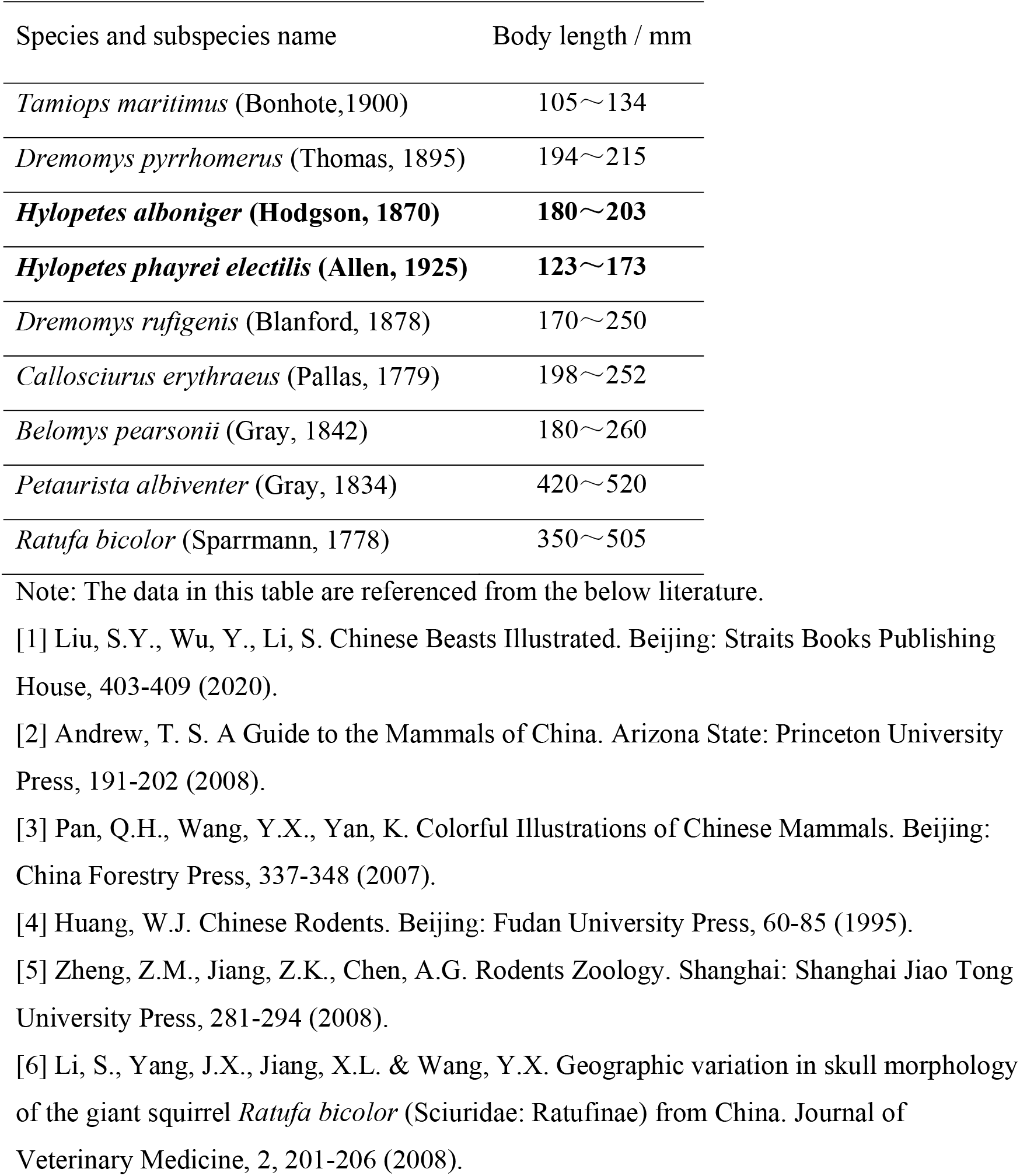
Nine squirrels living in Jianfengling, Hainan Island, China.

